# Altered astrocyte morphology in pathogenic *HEPACAM* variants: a multipoint analysis and new machine learning framework for 3D Sholl analysis

**DOI:** 10.64898/2026.09.02.748961

**Authors:** Madelyn G. Coble, Amy L. Stanek, Katherine T. Baldwin

## Abstract

Astrocytes are morphologically complex glial cells that play critical roles in brain development and function. Altered astrocyte morphology is associated with altered astrocyte function and is a common feature of many neurological disorders. Astrocytes express numerous membrane proteins that are important for their morphogenesis, including hepaCAM, an astrocyte-enriched cell adhesion molecule that regulates astrocyte branching organization, tiling, and coupling. Pathogenic variants of *HEPACAM* that impair homophilic protein interaction cause megalencephalic leukoencephalopathy with subcortical cysts (MLC), a rare and early-onset leukodystrophy characterized by white matter edema, seizures, and cognitive and motor decline. Pathogenic variants show altered subcellular localization and impaired interaction with key binding partners, but the impact on astrocyte morphology remains unexplored. Here we expressed three different dominant pathogenic *HEPACAM* variants in astrocytes of the developing mouse cortex and performed a comprehensive multipoint analysis of astrocyte morphology. Using established analysis workflows and a new machine learning model for efficient 3D Sholl analysis, we found small, but significant increases in morphological complexity for the G89S pathogenic variant, which impairs homophilic *cis* interaction of hepaCAM, and the Q56P pathogenic variant, which impairs homophilic *trans* interaction. This phenotype is distinct from the morphological changes we previously observed in *Hepacam* knockout astrocytes, suggesting that dominant variants may impact astrocyte morphology through a gain-of-function, rather than a loss-of-function, mechanism. Our study also provides a new machine learning workflow for streamlined 3D analysis of astrocyte branching complexity along with a framework for performing multivariate analysis of astrocyte morphology metrics across multiple conditions.

## Introduction

Astrocytes are structurally and functionally complex glial cells that are ubiquitous throughout the brain. Astrocytes interact with the various cells and structures in their microenvironment, via their elaborately branched arbors, to regulate critical brain functions such as synapse formation, ion and water homeostasis, and neurovascular coupling. Altered astrocyte morphology is associated with changes in gene expression and functional state, impaired cell-cell communication, and altered neuronal activity (Baldwin et al., 2023; Deng et al., 2026; Endo et al., 2022; Stogsdill et al., 2017). Further, altered astrocyte morphology is a hallmark feature of numerous neurological disorders (Farina et al., 2023; Hayatdavoudi et al., 2022; Schiweck et al., 2018).

Hepatic and glial cell adhesion molecule (hepaCAM, also known as GlialCAM) is an abundant, astrocyte-enriched, transmembrane protein that plays multiple important roles in astrocyte morphogenesis. In the developing mouse cortex, hepaCAM regulates astrocyte branching organization, territory establishment, gap junction coupling, endfoot development, synaptic strength, and neurite outgrowth (Baldwin et al., 2021; Gilbert et al., 2019; Jin et al., 2023). Missense mutations in human *HEPACAM* cause megalencephalic leukoencephalopathy with subcortical cysts (MLC), an early-onset and slowly progressive leukodystrophy characterized by macrocephaly, white matter edema, seizures, and motor and cognitive decline (Hamilton et al., 2018; Lopez-Hernandez et al., 2011; Pla-Casillanis et al., 2022). Disease-causing mutations, hereafter referred to as pathogenic variants, may be dominant or recessive depending on the specific amino acid substitution, leading to distinct MLC phenotypes.

Heterozygous dominant variants cause MLC Type 2b, a less severe form of the disease with a higher instance of autism spectrum disorder (Bosch & Estevez, 2020; Hamilton et al., 2018; Lopez-Hernandez et al., 2011). Many leukodystrophies, including MLC, Alexander Disease, and Vanishing White Matter Disease, originate from genetic mutations that disproportionately impact astrocyte function (Brenner et al., 2001; Jorge & Bugiani, 2019; Kerst et al., 2025; Trinanes-Ramos et al., 2025) yet few studies have examined the impact to astrocyte structure.

In a recent study, we examined the impact of three different dominant pathogenic variants on hepaCAM protein localization and interactome (Lewis et al., 2026). In mouse cortical astrocytes, endogenous mouse hepaCAM and exogenously expressed human hepaCAM display a highly regulated and punctate distribution pattern throughout the astrocyte membrane, including at cell-cell junctions and endfeet (Baldwin et al., 2021). In contrast, exogenously expressed dominant pathogenic variants G89S, Q56P, and D128N showed substantially altered subcellular distribution, with diffuse expression homogeneously distributed throughout the cell (Lewis et al., 2026). HepaCAM interacts with itself both in *cis* and in *trans* via its N-terminal IgV domain. Most pathogenic variants, including the ones we tested, occur in the IgV domain. G89S is located at site of homophilic *cis* interaction, and has been shown to disrupt both homophilic *cis* and *trans* interaction, suggesting that *cis* interaction is a prerequisite for *trans* interaction (Elorza-Vidal et al., 2020). Q56P and D128N are found at the *trans* interaction site and impair *trans* interaction, leaving *cis* interaction intact (Elorza-Vidal et al., 2020). Whether pathogenic variants impact astrocyte morphology *in vivo*, and whether this impact differs between variants has not been explored.

Here we used an established viral strategy to express dominant pathogenic *HEPACAM* variants in astrocytes of the developing mouse cortex and performed a comprehensive multipoint assessment of astrocyte morphology. In addition to using established methods for analyzing astrocyte morphology, we developed a trained model for efficient 3D Sholl analysis of <u>d</u>econvolved *in <u>v</u>ivo* <u>a</u>strocytes (DIVA) using Imaris AI-Filament Tracer along with a user-friendly script for multivariate analysis of astrocyte morphology.

## Results

### Viral expression of exogenous hepaCAM in early postnatal development

To investigate the impact of dominant pathogenic *HEPACAM* variants on astrocyte morphology in the developing mouse brain, we used a previously established astrocyte-specific viral expression system (Baldwin et al., 2021; Lewis et al., 2026). Plasmids containing wild-type (WT) human *HEPACAM* cDNA or one of three different pathogenic variant human *HEPACAM* cDNAs with a C-terminal TurboID enzyme and HA tag under control of the human minimal GFAP promoter (gfaABC1D) were packaged into PHP.eB adeno-associated virus (AAV) and administered via unilateral intracortical injection to neonatal CD1 mice between postnatal day 1 (P1) and P2 (**Figure 1A**). For morphological investigation, we chose three different dominant variants that cause MLC Type 2b in the presence of one normal *HEPACAM* allele: G89S which impairs *cis* interaction, Q56P which impairs *trans* interaction, and D128N which impairs *trans* interaction and introduces a new glycosylation site (Elorza-Vidal et al., 2020). To visualize astrocyte morphology, mice were administered a second AAV to express membrane-targeted mCherry-CAAX. To control for any changes in morphology that might occur due to overexpression of hepaCAM protein, we generated a control group that was administered only the mCherry-CAAX virus. We previously validated the efficiency and specificity of AAV-mediated astrocyte-specific expression of WT and variant hepaCAM protein the mouse cortex at P21 (Lewis et al., 2026). We also previously demonstrated that inclusion of a C-terminal TurboID enzyme does not alter the subcellular distribution of hepaCAM, nor does it impact its association with known interaction partners (Lewis et al., 2026).

**Figure 1:**
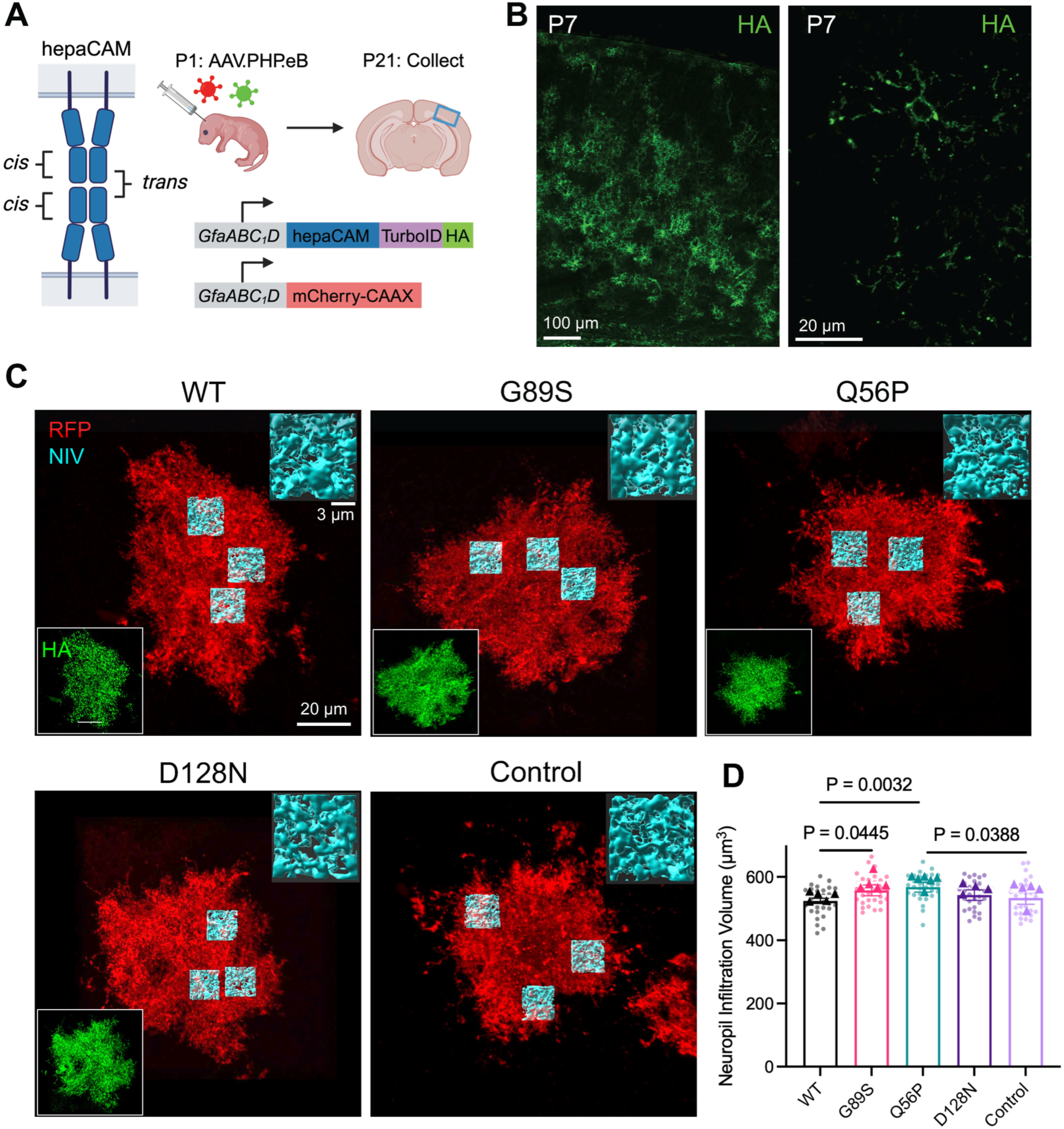
Impairing homophilic *cis* and *trans* binding in hepaCAM increases astrocyte neuropil infiltration. **A)** HepaCAM interacts with itself in *cis* and in *trans* at cell-cell junctions. P1 mice were administered intracortical AAVs to express hepaCAM-Turbo-HA fusion proteins and mCherry-CAAX. Brains were collected at P21 for imaging of astrocytes in layer 5 of the visual cortex (VCX, blue box) followed by morphology analysis. Created with BioRender.com. **B)** Left: tile scan image and right: high magnification image of mouse VCX at P7 showing robust expression of WT hepaCAM-Turbo-HA (green). **C)** Representative images of VCX L5 astrocytes at P21 expressing mCherry-CAAX (RFP, red) and hepaCAM constructs (WT, G89S, Q56P, D128N) and mCherry-CAAX control astrocytes. HA expression (green) is shown in the inset on the bottom left. Scale bar 20 µm. Neuropil infiltration volume (NIV, cyan) reconstructions are shown in the top right for each image. **D)** NIV analysis. Data presented as animal averages (represented as triangles; n = 4–6 mice per group) and individual astrocytes (represented as circles; n = 30 cells per condition), bars are mean ± SEM. Normality determined using Shapiro-Wilk test. P-values computed with one way ANOVA with Tukey’s Posttest.

In protoplasmic astrocytes of the mouse cortex, astrocyte morphogenesis occurs postnatally, with astrocytes migrating and establishing major branches during the first postnatal week and growing in size and complexity during the second and third postnatal weeks (Stogsdill et al., 2017; Watanabe et al., 2023). HepaCAM protein expression in the mouse cortex is low at P1 and increases substantially by P7, remaining high throughout life (Baldwin et al., 2021). To confirm that our viral strategy resulted in exogenous hepaCAM expression by the end of the first postnatal week, we collected brains from AAV-injected mice at P7 and performed HA staining to detect exogenous hepaCAM expression. We observed robust expression of hepaCAM-HA throughout astrocyte branches at P7 (**Figure 1B**), indicating that exogenous proteins are present during the second and third postnatal weeks, when astrocytes undergo extensive elaboration of fine branches.

### Astrocytes expressing G89S and Q56P variants show increased neuropil infiltration volume

To assess the cell-autonomous impact of dominant pathogenic variants on astrocyte morphology, we collected brains from P21 mice and acquired high resolution confocal images of individually transduced astrocytes from layer 5 of the mouse visual cortex. We focused our studies on layer 5 astrocytes in the visual cortex because astrocyte morphogenesis is well characterized in this region (Stogsdill et al., 2017; Watanabe et al., 2023), and we have previously investigated the impact of *Hepacam* deletion on astrocyte morphology in this region (Baldwin et al., 2021). We first investigated whether pathogenic variants impact the ability of astrocytes to form fine branching structures in the synapse-rich neuropil by assessing astrocyte neuropil infiltration volume (NIV) (**Figure 1C**). We did not observe any difference in NIV between astrocytes transduced with WT hepaCAM (hereafter referred to as WT) and control astrocytes (**Figure 1D**), indicating that WT hepaCAM overexpression does not significantly impact astrocyte complexity. Both G89S and Q56P pathogenic variants showed significantly increased NIV compared to WT; however, we did not observe any significant differences in NIV for D128N variants. Collectively, these results suggest that both *cis* and *trans* impairing variants of hepaCAM increase the extent to which an astrocyte infiltrates its surrounding environment, but there may be differences in the potency of specific pathogenic variants.

### Territory and surface volumes are increased in G89S mutant astrocytes

Astrocytes tile the brain, forming distinct, non-overlapping territories(Bushong et al., 2002). Previous research has shown that sparse deletion of *Hepacam* reduces astrocyte territory *in vivo* and impairs astrocyte tiling behavior (Baldwin et al., 2021). Therefore, we next sought to determine whether pathogenic variants of *HEPACAM* alter astrocyte cell volume or territory volume. To do so, we acquired images of complete transduced astrocytes contained within 100 µm thick tissue sections (**Figure 2A**). Incomplete astrocytes were excluded from analysis. Using our established Imaris workflows, we calculated astrocyte cell volume, territory volume, and relative complexity (cell volume divided by territory volume) (**Figure 2B**). We did not observe any difference between WT and control territory volume in any of the measurements, indicating that WT overexpression does not significantly alter astrocyte territory volume. We observed that G89S-expressing astrocytes had significantly larger cell and territory volumes than control astrocytes (**Figure 2C-D**). Combined with the initial findings from NIV analysis, this highlights a potential unique role for hepaCAM *cis* interactions in astrocyte morphogenesis.

**Figure 2:**
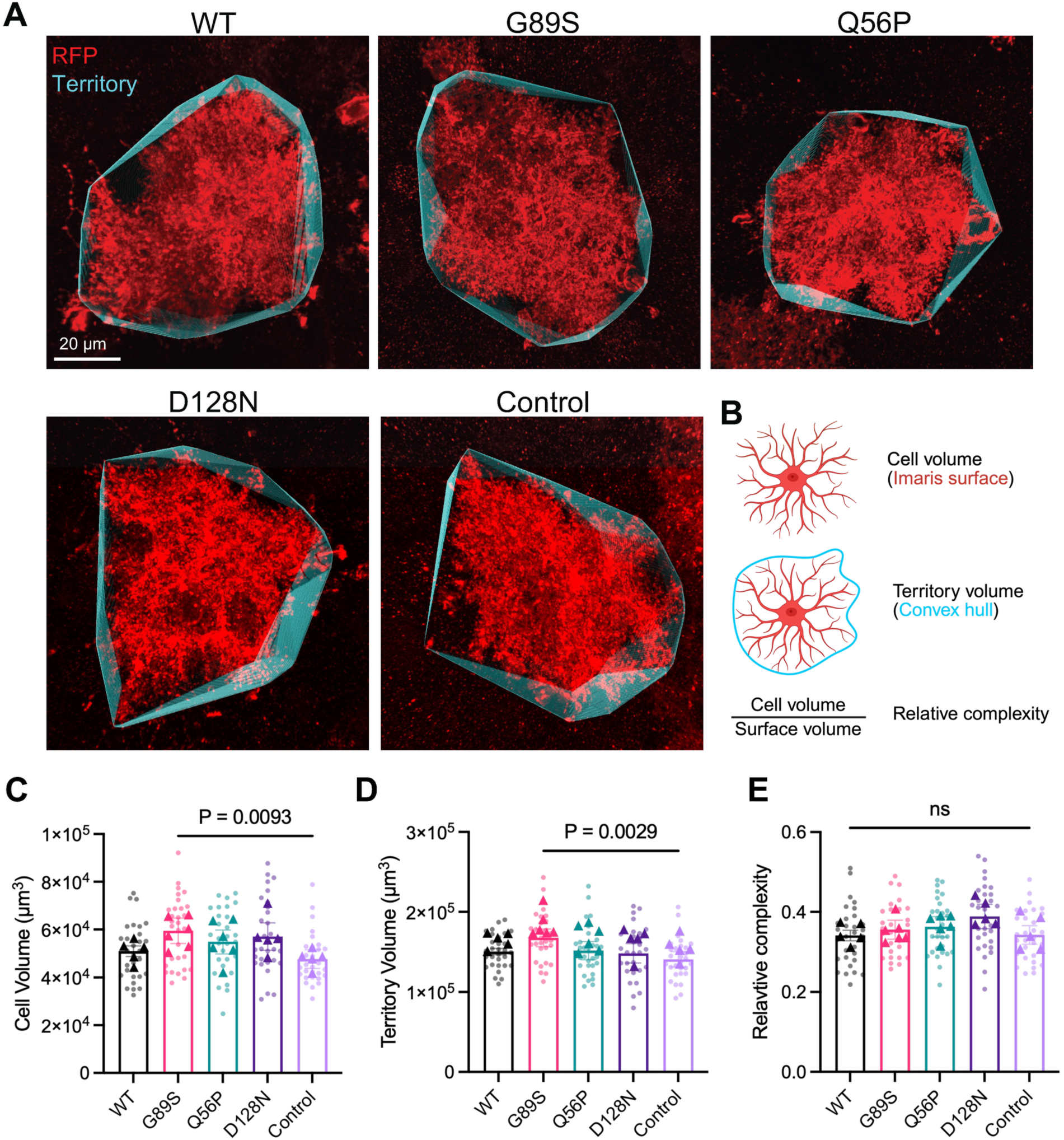
G89S mutant astrocytes with impaired *cis* homophilic binding have increased surface and territory volumes. **A)** Representative images of VCX L5 astrocytes expressing mCherry-CAAX (RFP, red) and hepaCAM constructs (WT, G89S, Q56P, D128N) and mCherry-CAAX control astrocytes with convex hull (territory) in cyan. **B)** Overview of morphology metrics. Cell volume: the volume of the 3D Imaris surface representing the fluorescent signal. Territory volume: the volume of the convex hull containing the surface. Relative complexity: calculated by dividing cell volume by surface volume. **C–E)** Whole-cell morphology analysis. Created with BioRender.com. **C)** astrocyte cell surface volume, **D)** astrocyte territory volume, and **E)** relative complexity. Data presented as animal averages (represented as triangles; n = 5–6 mice per group) and individual astrocytes (represented as circles; n = 30 cells per condition), bars are mean ± SEM. Normality determined using Shapiro-Wilk test. P-values computed with **C)** Kruskal-Wallis Test with Dunn’s Posttest and **D-E)** one-way ANOVA with Tukey’s Posttest.

### Development of a trained model for efficient analysis of astrocyte branching complexity

While both NIV and territory analysis produce useful information about astrocyte complexity and size, neither measurement captures the overall complexity of astrocyte branching architecture. 3D Sholl analysis has been previously used to assess branching complexity of *in vivo* astrocytes but, due to the structural complexity of astrocytes, requires a significant time commitment and involves extensive manual tracing and correction to generate accurate filament traces of astrocytes (Tan et al., 2023; Testen et al., 2025). To overcome this barrier, we used the AI filament tracer feature in Imaris 10 to develop an optimized workflow for analysis of astrocyte branching complexity *in vivo*, reducing the time spent analyzing each cell to only a few minutes. We used a data set of 100 deconvolved images of complete mCherry-CAAX transduced astrocytes with or without exogenous hepaCAM to train the AI filament tracer to produce 3D filament reconstructions of astrocytes suitable for 3D Sholl analysis (**Figure 3**). We refer to our trained model as DIVA (<u>d</u>econvolved *in <u>v</u>ivo* <u>a</u>strocyte) filament. Using DIVA filament followed by 3D Sholl analysis, we observed a peak Sholl complexity of 300 intersections at approximately 20 µm from the soma for WT and control astrocytes, similar to other studies that performed 3D Sholl analysis of cortical astrocytes at P21 (Sakers et al., 2026; Tan et al., 2023) (**Figure 4A, B**). 3D Sholl analysis of astrocytes expressing WT or pathogenic variants did not reveal any significant differences in complexity of the branching architecture (**Figure 4B**), nor was there a difference in the average number of total interactions (Figure 4C). We did observe a significant increase in peak complexity in the G89S condition compared to D128N, but we did not observe significant differences between pathogenic variants and WT or control (**Figure 4D**). Overall, this result indicates that, despite increased NIV in G89S and Q56P conditions, 3D branching architecture is not significantly different between WT and pathogenic variants.

**Figure 3:**
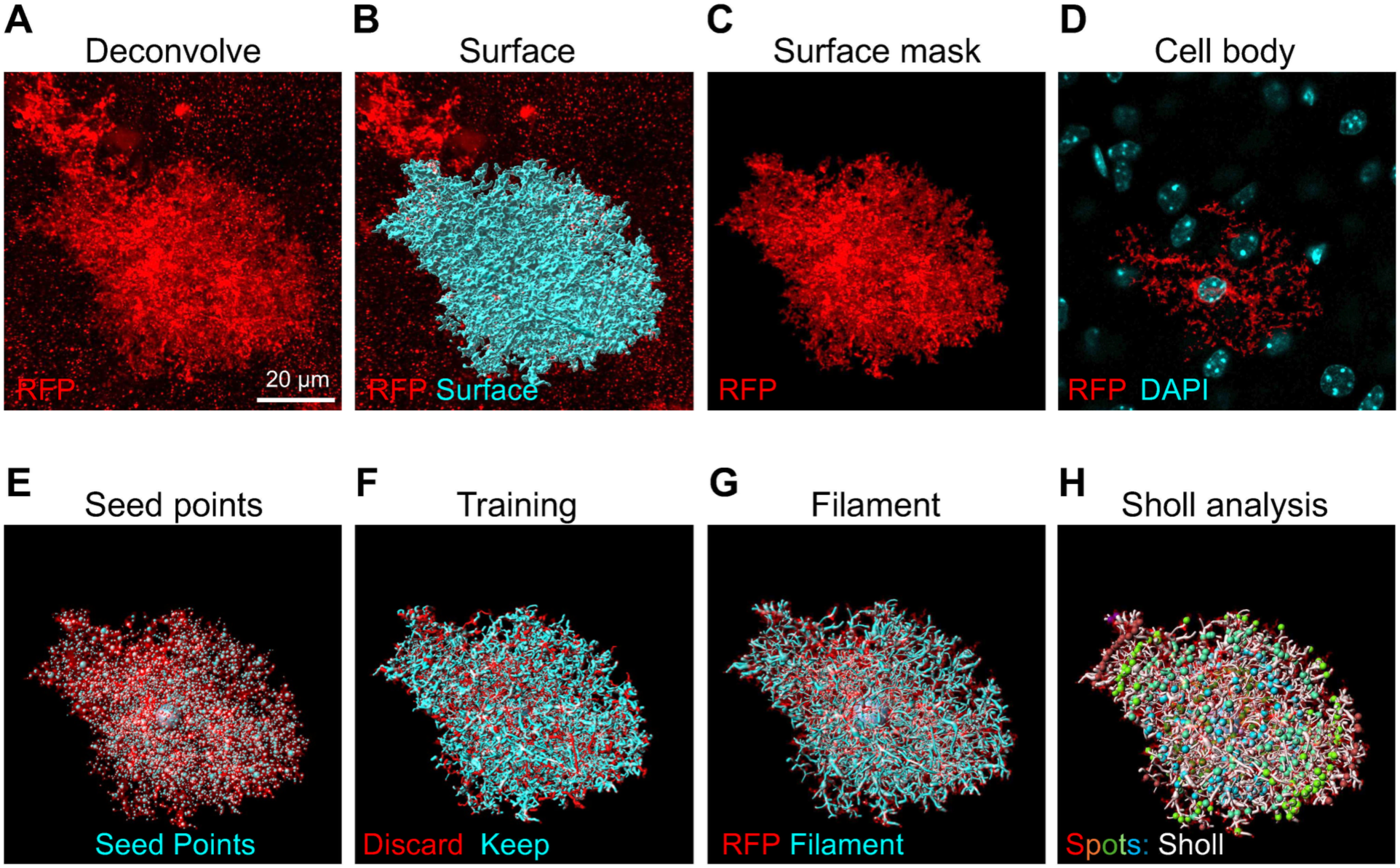
DIVA (<u>D</u>econvolved *In <u>V</u>ivo* <u>A</u>strocyte) filament analysis in Imaris. **A)** Deconvolved image in 3D view, mCherry-CAAX (RFP) in red. Scale bar 20 µm. **B)** Creation of stringent surface (cyan) from mCherry-CAAX (red) labeling. **C)** Surface mask applied to isolate the mCherry-CAAX (red) signal within the surface. **D)** Identification of astrocyte cell body using DAPI staining (cyan) to serve as starting point for filament construction. **E)** Use of multiscale seed points (cyan) to establish branching path. **F)** Use of machine learning segment classification to train DIVA AI and construct the filament, “discard” filaments in red, “keep” filaments in cyan. **G)** Finalized filament (cyan), visualized with mCherry-CAAX channel for manual check for filament construction errors. **H)** Use of adapted MATLAB Filament Sholl Analysis XTension to detect Sholl intersections (multicolored spots).

**Figure 4:**
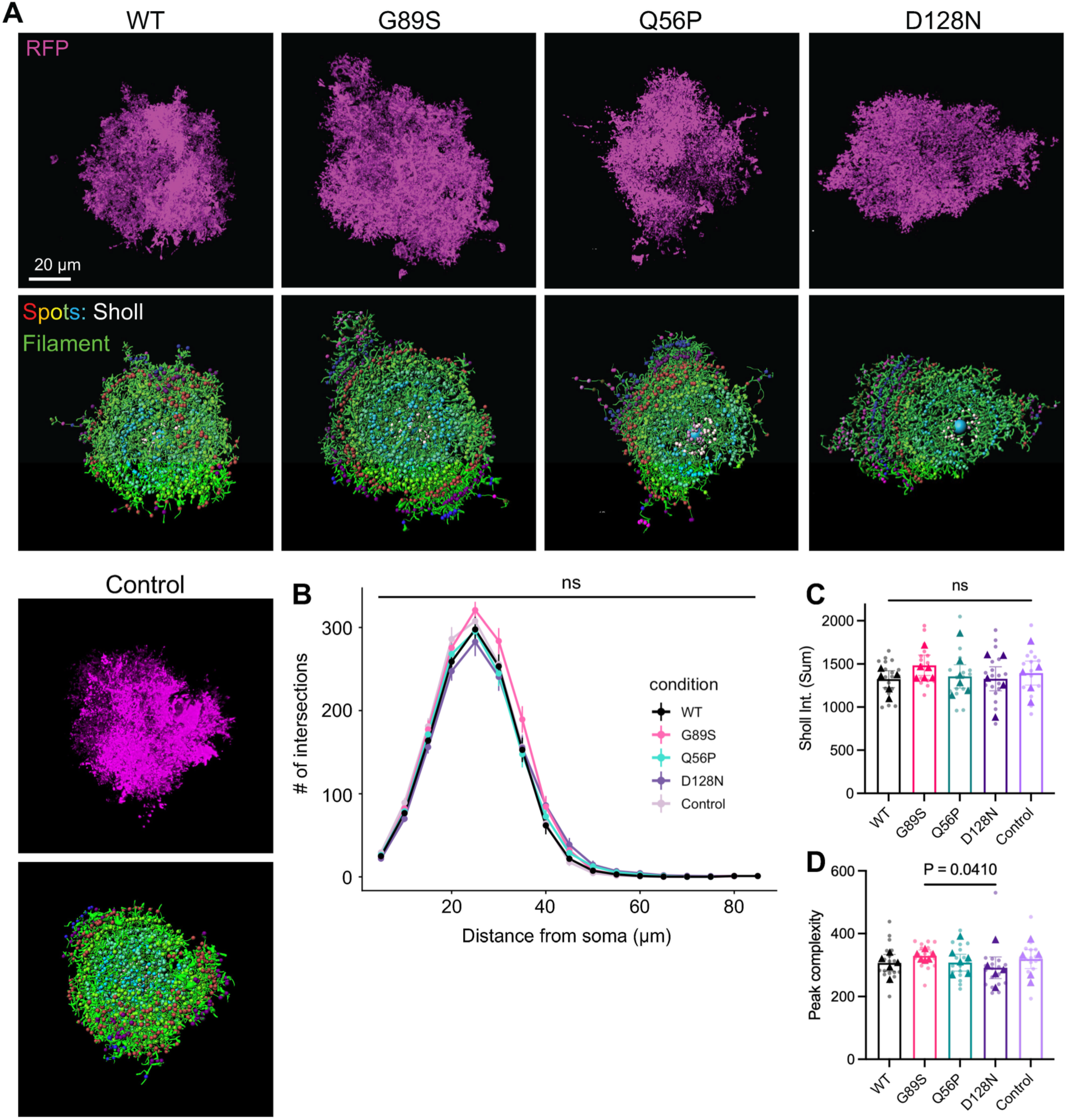
Branching complexity is not significantly altered by pathogenic variant expression. **A)** Representative images of VCX L5 astrocytes expressing deconvolved mCherry-CAAX (top, magenta) and constructed DIVA filament (bottom, green). Sholl intersections represented as multicolored spots). **B)** 3D Sholl analysis curves for *in vivo* hepaCAM overexpression (WT, G89S, Q56P, D128N) and mCherry-CAAX control astrocyte branching complexity. n = 5–6 mice per group, n = 16–19 cells per condition. Data were analyzed using a mixed effects model in RStudio, presented as mean ± SEM. Significance of fixed effects was assessed using ANOVA in the mixed model with Tukey’s Posttest. **C)** Quantification of total Sholl intersections per condition. **D)** Quantification peak complexity (Sholl curve peak). **C-D)** Data presented as animal averages (represented as triangles) and individual astrocytes (represented as circles), bars are mean ± SEM. Normality determined using Shapiro-Wilk test. P-values computed with **C)** one way ANOVA with Tukey’s Posttest and **D)** Kruskal-Wallis Test with Dunn’s Posttest.

### Multivariate analysis of astrocyte morphology across conditions

The common strategies for analyzing astrocyte morphology via fluorescence microscopy that we have used in this manuscript all offer important insight into astrocyte structure; however, each is limited in its ability to provide a complete assessment of astrocyte morphology. For example, 3D Sholl analysis reveals overall branching architecture, yet struggles to resolve the finer details visible with NIV analysis. Multivariate image analysis is a useful tool for simultaneously visualizing overall impact of different conditions (e.g., pathogenic variants) on multiple parameters and has been recommended as a method for analyzing astrocyte morphological complexity (Barriola et al., 2025). We developed an annotated R-script to perform multivariate analysis of astrocyte morphology metrics across multiple conditions. Using this pipeline, we performed principal component analysis (PCA) using six morphological measurements: NIV, territory volume, surface volume, relative complexity, number of Sholl intersections (intersections), and peak of Sholl curve (peak). We visualized the results as a score plot, with subjects represented as individual points colored by experimental condition and 95% confidence ellipses plotted around each group centroid (**Figure 5A**). We also generated a loading plot (**Figure 5B**) and a biplot (**Figure 5C**) to visualize the contribution of individual morphological variables to each principal component. Finally, we performed statistical analysis to determine whether there was any significant difference between conditions. Uncorrected p-values showed a significant difference (p < 0.05) between WT and G89S and control and G89S, though correction for multiple comparisons pushed the p-value above 0.05 (**Figure 5D**). Overall, these results are consistent with our findings that expression of G89S in developing astrocytes significantly impacts some morphological parameters and not others.

**Figure 5:**
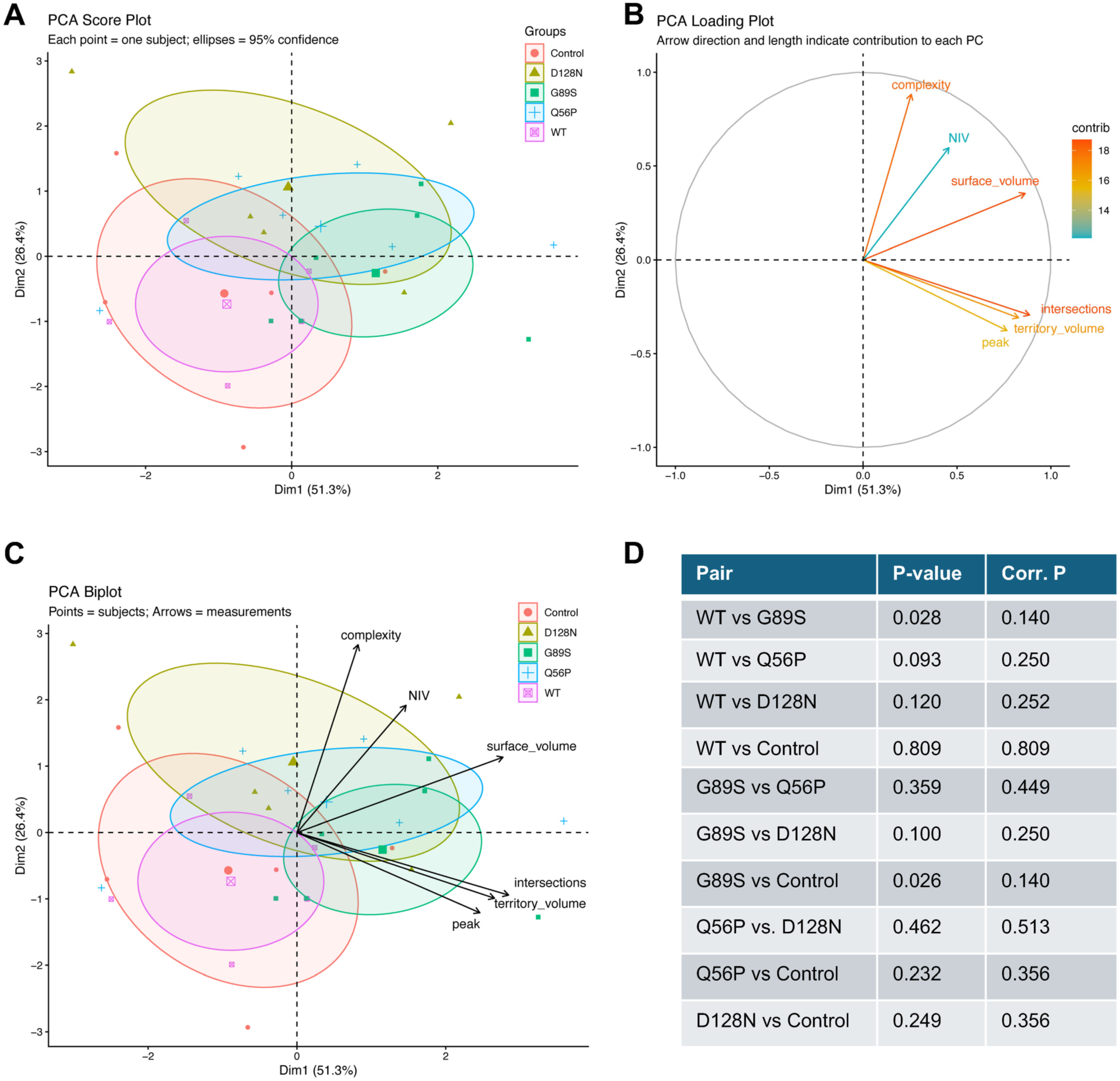
Multivariate analysis of astrocyte morphology. **A)** PCA Score Plot with each subject represented by one point. Ellipses represent 95% confidence intervals. **B)** PCA Loading Plot with arrow direction and length indicating the contribution of each measurement to each principal component. **C)** A PCA Biplot, showing information from the score plot with arrows from the loading plot overlayed. **D)** Table showing the uncorrected P values (P-value) and corrected P values (Corr. P) for each individual comparison following PERMANOVA analysis.

## Discussion

Here we investigated the impact of pathogenic hepaCAM variants on astrocyte morphology in the developing mouse cortex. We used a previously established viral expression strategy to introduce WT hepaCAM or one of three different dominant pathogenic variants into astrocytes at P1. At P21, we assessed multiple morphological parameters and found increased NIV in G89S and Q56P variants, as well as increased territory volume in G89S. Moreover, we developed new methods for analyzing astrocyte morphology, including a trained model for generating astrocyte filaments and a pipeline for multivariate analysis.

Of the three pathogenic variants that we tested, G89S had the strongest effect on astrocyte morphology. Both G89S and Q56P increased astrocyte NIV, but only G89S increased astrocyte territory volume. Moreover, G89S was the only condition to show significant difference from WT and control conditions in the multivariate analysis, albeit with uncorrected p-values.

Previous studies have shown that G89S mutation occurs in the *cis* binding site and impairs both homophilic *cis* and *trans* interaction. In contrast, Q56P and D128N are located at *trans* interaction sites and impair only *trans* interaction (Elorza-Vidal et al., 2020). Because all pathogenic variants lack *trans* interaction, but only G89S lacks *cis* interaction, our results point to a potential unique role for hepaCAM *cis* interaction in regulation of astrocyte morphology. We previously found that *Hepacam* knockdown in isolated astrocytes reduces astrocyte territory volume, with no significant impact to NIV (Baldwin et al., 2021). Thus, our finding that G89S increases territory volume and NIV suggests that this pathogenic variant confers gain-of-function with respect to astrocyte morphology, rather than operating as a dominant negative.

This does not preclude the possibility that pathogenic variants also cause loss of normal hepaCAM function. Indeed, our recent comparison of WT and pathogenic variant interactomes reveals both decreased association with key transmembrane proteins and increased or new association with others (Lewis et al., 2026), supporting the idea that pathogenic variants may cause both gain and loss of function. While many changes we observed in our proteomic dataset were common to all three variants, some were unique to each variant. These unique targets may provide mechanistic insight into the differing impact of *cis* versus *trans* mutations on hepaCAM function in future studies.

Further investigation is needed to understand the biological importance of increased NIV in the G89S and Q56P conditions. Previous studies have detected significant differences in astrocyte NIV with corresponding changes in synaptic activity. For example, deletion of *Nlgn2* from astrocytes significantly reduces NIV and decreases excitatory synapse number and excitatory synaptic transmission, while increasing inhibitory transmission (Stogsdill et al., 2017). Conversely, deletion of *Nrcam* from astrocytes increases NIV with decreases in inhibitory synapse number and inhibitory transmission (Takano et al., 2020). The full picture is more complex, as deletion of *Hepacam* from astrocytes causes no observable change in NIV yet significantly increases excitatory synaptic strength and decreases inhibitory synaptic strength (Baldwin et al., 2021). Whether the increased NIV in the pathogenic variant conditions is accompanied by changes in astrocyte-synapse association and/or synaptic function is a topic that requires further investigation.

Analysis of astrocyte NIV in thinner tissue sections (40 µm) enables use of higher magnification objectives and avoids issues with antibody and laser penetration, enabling detection of more and finer astrocyte processes than is typically possible in thick tissue sections. However, NIV is an imperfect measurement as it only samples regions of neuropil within the astrocyte domain and does not capture entire cells. We therefore expanded our analysis to include full astrocytes. Though we detected increased NIV for G89S and Q56P variants, we did not detect any significant changes in relative complexity of full astrocytes or in astrocyte branching complexity via 3D Sholl analysis. This discrepancy is likely a result of the many technical challenges associated with analysis of complete astrocyte morphology.

Capturing the entire cell volume of membrane-labeled protoplasmic astrocytes at a high enough resolution to detect fine leaflet processes remains a challenge within the field. Reliable capture of entire astrocytes in the z-dimension requires thick 80 – 100 µm tissue sections. Challenges with antibody penetration, objective working distance, poor resolution in the z-dimension, and laser penetration beyond 100 µm all impact the ability to detect finer astrocytes processes.

Moreover, many fine processes are beyond the resolution limit of light microscopy and require super resolution or electron microscopy for detection (Baldwin et al., 2023; Salmon et al., 2023). As a result, many of these fine details are filtered out during creation of surfaces and filaments in Imaris, leading to an artificial reduction in astrocyte complexity. Nonetheless, the trained DIVA filament construction method that we created will serve as a powerful tool for the research community, dramatically improving the efficiency of analyzing the complex branching architecture of protoplasmic astrocytes. Combining DIVA filament analysis with labeling strategies that have proven highly effective for visualizing small astrocyte processes, such as spaghetti monster fluorescent proteins (Gleichman et al., 2025), could further expand its utility.

As highlighted above, the many nuances of astrocyte morphology cannot be captured in a single morphological parameter, such as territory volume, NIV, or Sholl analysis. As the number of potentially informative morphology metrics increases, there is an apparent need for comprehensive tools to make sense of the growing pile of data. Multivariate image analysis is one useful strategy for meeting this need. We therefore developed a user-friendly R-script for performing PCA analysis of multiple morphological measurements and applied this to our data. Visualization on a PCA Score Plot showed a strong spatial relationship between the WT and control conditions, demonstrating the robustness of our approach. All three pathogenic variants were shifted rightward and upward compared to WT and control conditions, and G89S was significantly different from WT and control conditions prior to correction for multiple comparisons. The Loading Plot and Biplot show the contribution of the different measurements to each principal component. In data sets with strong morphological differences between conditions, these plots could provide useful insight into the relationship between different morphological parameters.

There are limitations to our study that should be considered when interpreting the data. The use of a viral over expression system to express the pathogenic variants creates unphysiologically high levels of hepaCAM protein expression. In our previous study, we showed that overexpressed wild type hepaCAM displays a similar localization pattern and puncta density to endogenous hepaCAM, with increased puncta intensity indicating overall higher expression levels (Lewis et al., 2026). Our overexpression constructs also contain a C-terminal fusion to the TurboID enzyme. This fusion protein was created for use in our previous proteomic study and was shown to have no impact on hepaCAM protein localization or expression.

Moreover, these fusion proteins showed robust interaction with known hepaCAM interaction partners, indicating that the TurboID enzyme dose not impact hepaCAM function (Lewis et al., 2026). Importantly, we compared all of our pathogenic variant conditions to the WT overexpression condition with C-terminal TurboID fusion. We also found no differences in astrocyte morphology between the WT overexpression condition and the control astrocytes expressing only mCherry-CAAX. We also focused our studies exclusively on dominant pathogenic variants, as these variants cause MLC Type 2b in humans in the presence of a wild-type *HEPACAM* allele. Still, the unphysiologically high levels of hepaCAM protein could have unanticipated impacts to astrocyte function and follow-up experiments with physiological levels of pathogenic variants will serve as important validation of these findings.

Additionally, the mosaic expression of viral constructs should be considered when interpreting the data. As demonstrated in our prior study, our neonatal AAV administration strategy is effective in transducing deeper layer astrocytes in the mouse cortex, producing both isolated transduced astrocytes and areas of contiguous transduced astrocytes (Lewis et al., 2026). For detailed morphological analysis, individually labelled astrocytes are necessary, as the complexity of astrocytes makes it extremely challenging to distinguish the territories of neighboring astrocytes that express the same fluorescent label. Thus, although our approach enables investigation of cell-autonomous function of pathogenic variants surrounded by non-transduced astrocytes, it does not enable investigation of how pathogenic variants impact astrocyte morphology at the population level. Given the known roles of hepaCAM in astrocyte territory establishment and gap junction coupling, future studies investigating the role of pathogenic variants in contiguous astrocytes, perhaps using a mosaic labeling system, will shed light on the role of pathogenic variants in astrocyte competition for territory and tiling.

Overall, our findings provide insight into the impact of pathogenic variants of *HEPACAM* and impaired *cis* interaction on astrocyte morphology. This advances our understanding of mechanisms that regulate astrocyte morphology in both physiological and pathological conditions. Moreover, our study provides the field with useful tools to improve the efficiency and effectiveness of astrocyte morphological analysis.

## Author Contributions

M.G.C: conceptualization, methodology, investigation, formal analysis, visualization, writing – original draft. A.L.S.: investigation, visualization. K.T.B: conceptualization, methodology, investigation, formal analysis, supervision, funding acquisition, visualization, writing – original draft.

## Acknowledgments

We thank Deniz Kesman for assistance with sample collection and sample processing.

## Materials and Methods

### Animals

All mice were used in accordance with the Institutional Animal Care and Use Committee (IACUC) and the UNC Department of Comparative Medicine (IACUC Protocol Numbers 21-116.0 and 24-005.0). Mice were housed in standard conditions with 12-hour day/night cycles.

Timed-pregnant CD1 females were obtained from Charles River (RRFD: IMSR_CRL:022). Mice were used for experiments at postnatal day 21 (P21), or as specified in the text and figure legends. Mice of both sexes were included in all experiments.

### AAV production and administration

Plasmids with a pZac2.1 backbone expressing membrane targeted hepaCAM-TurboID-HA, G89S-TurboID-HA, Q56P-TurboID-HA, D128N-TurboID-HA, TurboID-HA, or mCherry-CAAX under control of the human minimal GFAP promoter (*GfaABC_1_D*) were generated as described previously (Lewis et al., 2026). Plasmids were packaged into AAV PHP.eB capsids by the UNC BRAIN Initiative Viral Vector Core as described previously (Lewis et al., 2026). CD1 mice aged P1–P2 were administered AAV via intracortical injection according to established lab protocols (Baldwin et al., 2021; Lewis et al., 2026). Briefly, mouse pups were anesthetized using hypothermia by incubating pups on ice for 3–5 minutes until cessation of movement and breathing was observed. All viruses were adjusted to the same titer (3.4×10^13^ GC/mL and 1 μL of AAV each virus (hepaCAM and mCherry-CAAX) was injected unilaterally into the cortex using a Hamilton syringe (701 RN) with a removable needle (Hamilton 7803–05 with a 33 gauge, point style 4, and length of 1.0 in).

### Immunohistochemistry

Mice were anesthetized with 0.8 mg/kg tribromoethanol (avertin) and perfused with 1x Tris Buffered Saline (TBS)/Heparin followed by ice cold 4% PFA in TBS. Brains were collected and post-fixed in 4% PFA overnight at 4°C. The following day, brains were washed 3x with TBS and transferred to 30% sucrose in TBS for cryoprotection at 4°C for 2 days. Brains were frozen in a medium containing 2 parts 30% sucrose in TBS and 1-part Optimal Cutting Temperature Compound (O.C.T.) and stored at −80°C until sectioning. Coronal sections containing the cortex were collected using CryoStar NX50 Cryostat (Thermo Fisher Scientific) alternating between two 100 μm slices and four 40 μm slices, and stored at −25°C in 50% glycerol in TBS.

Immunolabeling was performed as described previously (Eaker & Baldwin, 2022). Briefly, slices containing the visual cortex were washed 3x 10 minutes with TBS containing 0.2% Triton (TBST), incubated in blocking solution (TBST with 10% goat serum) for 1 hour at room temperature. Primary antibody solution was prepared by diluting anti-Rat HA (Sigma 11867423001, 1:500) and anti-Guinea Pig RFP (Synaptic Systems 390–004, 1:1000) in blocking solution and centrifuging for 5 minutes (4700 rpm, 4°C). Sections were incubated for 2 nights (40 µm) or 3 nights (100 µm) in primary antibody solution at 4°C. Following primary antibody incubation, slices were washed 3x 10 minutes with TBST. Secondary antibody solution was prepared by diluting Goat anti-Rat 488 (Invitrogen A–11006, 1:200) and Goat anti-Guinea Pig 594 (Invitrogen A–11076, 1:200) in blocking solution and centrifuging for 5 minutes (4700 rpm, 4°C). Slices were incubated in secondary antibody solution at room temperature for 2 hours (40 µm sections) or 3 hours (100 µm sections). Slices were then washed 3x 10 minutes with TBST. For 100 µm slices used for DIVA Filament Analysis, DAPI (1:50,000) was added to the first of the 3 washes. Following all incubations and washes, slices were mounted onto glass slides, excess liquid was aspirated from the slide, and a drop of homemade mounting media (90% glycerol and 0.5% N-propyl gallate in 20 mM Tris pH 8.0) was added to each brain slice. More mounting media was added for 100 µm brain slices (∼2 drops) to ensure preservation of thicker tissue sections. No 1.5 glass coverslips were applied, sealed with nail polish, and dried at room temperature for 30 minutes before storage at 4°C. Confocal images were collected within one week of staining to ensure maximum fluorescent output and to maintain high signal-to-noise ratio.

### Confocal Image Acquisition

For all morphology analysis, point scanning confocal images of transduced astrocytes co-expressing HA and mCherry-CAAX were collected in layer 5 of the visual cortex using an Evident Fluoview 3000RS confocal microscope, using 488 nm and 594 nm laser lines to excite HA and mCherry-CAAX, respectively. Laser power was adjusted manually for each cell to account for variability in viral transduction efficiency, while avoiding signal saturation. For 100 µm sections used for DIVA, a 405 nm laser was used to excite DAPI. Control astrocytes expressed mCherry-CAAX alone. Images for neuropil infiltration volume (NIV) analysis were collected from 40 µm sections using a 60x oil-immersion objective with 2x optical zoom (20–36 µm z-stack range, 0.50 μm step size); z-stack boundaries were set based on the presence of the selected astrocyte within the tissue section. Images of complete astrocytes for territory volume analysis were collected from 100 µm sections using a 40x oil-immersion objective with 2x optical zoom (80–120 µm z-stack range, 0.50 µm step size); z-stack boundaries were set based on the full extent of the astrocyte’s territory within the tissue section. All morphological analyses were performed using the mCherry-CAAX signal. Images at the P7 timepoint were acquired using a Leica Stellaris FALCON 8 confocal microscope with a 20x air objective (tiled image, 10 µm z-stack, 1 µm step size) or a 100x oil-immersion objective (5 µm z-stack, 0.3 µm step size).

### Neuropil Infiltration Volume (NIV) Analysis

Neuropil infiltration volume (NIV) analysis was completed as previously described (Eaker et al., 2026). Briefly, post-processing was completed in Imaris by applying the “Normalize Layers” filter to compensate for depth-dependent signal attenuation and to improve uniformity of fluorescence intensity across the z-stack. For each astrocyte, three ROIs (12.45 µm x 12.45 µm x 10 µm) containing only neuropil (excluding astrocyte soma, large branches, and endfeet) were selected and reconstructed in Imaris using the surface tool. Surface volume within individual ROIs was recorded and averaged for each cell (3 x ROIs) and animal (3 x ROIs across 4-5 cells per animal). Data were analyzed in GraphPad Prism 10 using a Shapiro-Wilk test to confirm normal distribution of data followed by a one-way ANOVA with Tukey’s post-test. The number of cells and animals analyzed is indicated in the corresponding figure legend for this experiment.

### Astrocyte Territory Analysis

Astrocyte territory analysis was completed as described previously with modifications (Eaker & Baldwin, 2022; Eaker et al., 2026). Inclusion criteria for analysis required the entirety of the astrocyte to be contained within a single brain section, specifically visual cortex layer 5. Astrocytes outside of this brain region and/or incomplete astrocytes were excluded from this study. Using Imaris, a surface was created based on mCherry-CAAX labeling that contained the entire cell, and thresholding was completed to ensure the constructed surface captured all fluorescently labeled astrocyte processes. Spots were generated close to the surface, and a custom MATLAB Convex Hull XTension file was used to create a convex hull close to the spots. The volume of the convex hull was recorded and reported as astrocyte territory volume. The volume of the surface was recorded and reported as cell volume. The relative complexity was calculated by dividing the cell volume by the astrocyte territory volume. Data were analyzed in GraphPad Prism 10 using a Shapiro-Wilk test to confirm normal distribution of data. Data that passed normality were analyzed using a one-way ANOVA followed by Tukey’s post-test. Data that failed normality were analyzed using the Kruskal-Wallis Test with Dunn’s multiple comparisons test. The number of cells and animals analyzed is indicated in the corresponding figure legend for this experiment.

### DIVA Filament Analysis

We developed a new method, DIVA Filament Analysis, to construct filaments using machine learning in Imaris. Prior to filament construction, 100 µm images of complete astrocytes were deconvolved using Olympus cellSens Imaging Software. The constrained iterative setting was used, selecting only the mCherry-CAAX channel for deconvolution and turning on background subtraction and noise reduction. To complete DIVA Filament analysis in Imaris, a refined surface was created of the deconvolved astrocyte, ensuring all branches and visible fine processes were included within the surface ROI with a fine surface detail of 1 pixel. Thresholding was adjusted to ensure the surface filled as much of the cell’s signal as possible without extending beyond its boundaries, and the number of voxels per image was set to 1. The lasso tool was used to manually delete erroneous surface pieces created, and this completed surface was used to create a masked mCherry-CAAX channel of the ROI for filament tracing.

With the deconvolved masked channel selected, the tree auto path feature was used for filament creation. The astrocyte cell body was located using the slicer view, and DAPI staining was used to verify the position of the cell body. This cell body was measured (6.5–9.0 µm in diameter), and a starting point was manually added to the structure. Multiscale seed points were calculated by using the slicer view to locate the smallest filament (0.22–0.45 µm wide) and the largest filament (1.20–2.20 µm wide). As astrocytes exhibit substantial heterogeneity in soma size and in branching diameters, these parameters were manually optimized for each cell. The seed point threshold was adjusted to its maximum value to ensure maximal seed point detection.

Machine learning for segment classification was used to train the Imaris AI. This iterative process was completed by training one 2D section of a cell at a time using the slicer view, and multiple segments at a time were selected as either ‘keep’ or ‘discard’ elements. Training took place until the desired filament was constructed, and these parameters were stored and updated with each subsequent filament construction. After finalizing the filament, the structure was manually checked for erroneous sections. Following the training process on 100 cells to generate the optimal DIVA AI, the dataset was re-analyzed using the same trained model to ensure consistency across measurements. To perform Sholl analysis, a MATLAB plugin from Imaris Open was adapted (Gastinger, 2018). This plugin found the number of Sholl intersections for a defined interval away from the soma and added a new Spots group for each interval exactly where the filament segment crossed the Sholl interval. Statistical analysis was completed in R using a mixed effects model, and the significance of fixed effects was assessed using ANOVA on the mixed model with Tukey’s Posttest (Tan et al., 2023). Data were visualized using RStudio.

### Multivariate Image Analysis

Multivariate analysis of astrocyte morphology was performed in R version 4.4.1 using principal component analysis (PCA). All six morphological measurements were z-score standardized (mean = 0, SD = 1) prior to analysis. PCA was carried out using the *prcomp* function with centering and scaling disabled to avoid double-standardization. Results were visualized as a score plot, with subjects represented as individual points colored by experimental condition and 95% confidence ellipses plotted around each group centroid. A loading plot and biplot were generated to examine the contribution of individual morphological variables to each principal component. All visualizations were produced using the factoextra package in R (v2.0.0).

To test whether experimental conditions differed significantly in multivariate morphological space, a permutational multivariate analysis of variance (PERMANOVA) was performed using the *adonis2* function from the vegan package (v2.7-3), based on a Euclidean distance matrix computed from the scaled measurements (999 permutations). Prior to interpreting the PERMANOVA result, homogeneity of within-group dispersion was assessed using the *betadisper* function followed by a permutation test (*permutest()*, 999 permutations). Where appropriate, pairwise post-hoc comparisons between conditions were conducted using the pairwiseAdonis package (v0.4.1) with Benjamini-Hochberg correction for multiple comparisons. An R script is available at https://github.com/BaldwinLabUNC/Astrocyte_morphology.

## Declaration of generative AI and AI-assisted technologies in the manuscript preparation process

During the preparation of this work the authors used Claude to assist in development of the R Script for multivariate image analysis. After using Claude to construct an initial framework, the authors carefully reviewed the code line by line and edited and annotated as needed for accuracy and clarity. The authors take full responsibility for the content of the published article.

## Ethics Statement

All experimental protocols were performed in accordance with NIH guidelines and received approval from the Animal Care and Use Committee of UNC Chapel Hill.

## Data Availability Statement

Custom analysis files and scripts are available https://github.com/BaldwinLabUNC/Astrocyte_morphology. Raw data files available upon request.

